# Nitric oxide negatively regulates gibberellin signaling to coordinate growth and salt tolerance in *Arabidopsis*

**DOI:** 10.1101/2022.02.13.480237

**Authors:** Lichao Chen, Shuhao Sun, Chun-Peng Song, Jian-Min Zhou, Jiayang Li, Jianru Zuo

## Abstract

In response to dynamically altered environments, plants must finely coordinate the balance between growth and stress responses for their survival. However, the underpinning regulatory mechanisms remain largely elusive. The phytohormone gibberellin promotes growth via a derepression mechanism by proteasomal degradation of the DELLA transcription repressors. Conversely, the stress-induced burst of nitric oxide (NO) enhances stress tolerance, largely relaying on NO-mediated *S*-nitrosylation, a redox-based posttranslational modification. Here, we show that *S*-nitrosylation of Cys-374 in the *Arabidopsis* RGA protein, a key member of DELLAs, inhibits its interaction with the F-box protein SLY1, thereby preventing its proteasomal degradation under salinity condition. The accumulation of RGA consequently retards growth but enhances salt tolerance. We propose that NO negatively regulates gibberellin signaling via *S*-nitrosylation of RGA to coordinate the balance of growth and stress responses when challenged by adverse environments.

## INTRODUCTION

To survive under fluctuating environments and unfavorable conditions, plants have evolved sophisticated mechanisms to cope with abiotic and biotic stresses (Zhou and Zhang, 2020; Zhu, 2016). Because available resources are limited and detrimental effects are imposed on plants by stress responses, tradeoff or fine-tuned balance between defense and growth is tightly controlled to allow better fitness for plants (Belda-Palazon et al., 2020; Smakowska et al., 2016; Verma et al., 2016; Yang et al., 2012). However, how plants coordinate growth and stress tolerance is poorly understood.

Phytohormones are key regulators modulating growth and stress tolerance in plants. Among those, gibberellin is a classic growth-promotion phytohormone that regulates a wide range of plant growth and developmental processes, including seed germination, root development, hypocotyl elongation, and flowering (Achard et al., 2007; Debeaujon and Koornneef, 2000; Huang et al., 2010; Shu et al., 2014; Ubeda-Tomas et al., 2008; Wilson et al., 1992). Gibberellin signaling is initiated by binding of the phytohormone to its receptor GA-INSENSITIVE DWARF1 (GID1). The activated GA-GID1 complex interacts with DELLAs to promote their association with the F-box protein SLEEPY1 (SLY1), eventually facilitating the proteasomal degradation of the repressor proteins (Daviere and Achard, 2013; Sun, 2011; Xu et al., 2014). The *Arabidopsis* genome contains a small gene family of 5 members encoding DELLA repressor proteins, namely GA-INSENSITIVE (GAI), REPRESSOR-OF-*ga1-3* (RGA), RGA-LIKE1 (RGL1), RGL2, and RGL3. Among those, *RGA* and *GAI* are major members of this small gene family, as mutations in these two repressors rescue the growth retardation phenotype of *sly1-10* mutant (Dill et al., 2004).

Extensive studies during the past decades have characterized DELLAs as key regulators of gibberellin signaling. DELLAs also act as links to connect with other signaling pathways. Notably, DELLAs physically interact with PHYTOCHROME-INTERACTING FACTOR, BRASSINAZOLE-RESISTANT1, and ETHYLENE INSENSITIVE3, to integrate signals of light and other phytohormones in coordinating plant growth (Achard et al., 2009; An et al., 2012; Bai et al., 2012; de Lucas et al., 2008; Feng et al., 2008; Ubeda-Tomas et al., 2009). In addition to the modulation of plant growth and development, DELLAs have also been found to regulate stress responses. While DELLA protein SiGAI4 positively regulates cold tolerance in tomato (Wang et al., 2020a) and the *Arabidopsis* gain-of-function mutant *gai-1* displays increased drought tolerance (Wang et al., 2020b), the *Arabidopsis gai-t6 rga-24* double mutant is sensitive to salt treatment (Achard et al., 2006). DELLA proteins also interact with JASMONATE-ZIM DOMAIN (JAZ) to retard the JAZ-MYC2 interaction, thereby enhancing the activity of MYC2 to modulate biotic stress responses (Hou et al., 2010).

Stress responses in plants are more often regulated by the stress-related phytohormones and other signaling molecules, including nitric oxide (NO). As an important signaling molecule, NO plays a vital role in regulating various physiological processes in all living organisms. In plants, NO regulates a wide range of biological processes, including flowering, reproductive development, seed germination, root and shoot development as well as responses to biotic and abiotic stresses (Duan et al., 2020; Fernandez-Marcos et al., 2011; He et al., 2004; Yu et al., 2014). The major bioactive NO species is *S*-nitrosoglutathione (GSNO) that is irreversibly degraded by the highly conserved GSNO reductase (GSNOR) (Liu et al., 2001). In *Arabidopsis*, mutations in the single-copied *GSNOR1* gene cause the accumulation of excessive amount of NO species, resulting in severe defects in development and stress responses (Chen et al., 2009; Feechan et al., 2005; Kwon et al., 2012; Lee et al., 2008). NO executes its physiological effects mainly through protein *S*-nitrosylation, a redox-based posttranslational modification by the addition of an NO molecule to the thiol group of cysteine residue (Cys) to form *S*-nitrosothiol (SNO) (Feng et al., 2019; Hess et al., 2005; Stamler et al., 1992). Protein *S*-nitrosylation modulates diverse functions of proteins, including enzymatic activities, subcellular localization, stability, and protein-protein interactions. In higher plants, mainly in *Arabidopsis*, a number of *S*-nitrosylated proteins have been reported to regulate various developmental processes, immune responses, stress responses, and phytohormone signaling (Astier et al., 2011; Feng et al., 2019; Yu et al., 2014).

The interplay between NO and phytohormone signaling has been studied in some degrees. In *Arabidopsis*, *S*-nitrosylation of the auxin receptor TIR1 enhances its interaction with the transcriptional repressors Aux/IAA to promote their proteasomal degradation (Terrile et al., 2012). In the cytokinin pathway, while *S*-nitrosylation of a histidine phosphotransfer protein negatively regulates the phosphorelay, leading to a compromised cytokinin response (Feng et al., 2013), NO chemically reacts with cytokinins to regulate the cellular homeostasis of NO (Liu et al., 2013), illustrating a fine-tuned reciprocal regulatory mechanism between these two classes of signaling molecules. In gibberellin signaling, the NO donor sodium nitroprusside (SNP) induces the accumulation of DELLAs (Lozano-Juste and Leon, 2011). Moreover, in response to environmental stress, *S*-nitrosylation of OST1 and ABI5 negatively modulates abscisic acid signaling (Albertos et al., 2015; Wang et al., 2015). These studies highlight the importance of NO-mediated *S*-nitrosylation in regulating both growth and stress responses in plants.

In spite of these efforts, the molecular mechanism regulating the balance between plant growth and stress responses remains largely elusive. In this study, we report that NO induces the *S*-nitrosylation of *Arabidopsis* DELLA protein RGA at Cys-374, which causes the inhibition of the RGA-SLY1 interaction, thereby stabilizing the RGA repressor protein to coordinate plant growth and abiotic stress responses.

## Results

### Nitric oxide negatively regulates gibberellin signaling via DELLA repressors

Gibberellin mainly promotes plant growth, a biological effect opposite to that of NO. To explore the possible interaction between the NO and gibberellin pathways, we tested the responses of *Arabidopsis* to these two signaling molecules. While gibberellin promoted the elongation of roots and hypocotyls, the NO donor sodium nitroprusside (SNP) inhibited the growth of roots and had no apparent effect on the elongation of hypocotyls (Figure 1A and 1B). The lack of inhibitory effect on hypocotyl elongation is likely attributed to the relatively low concentrations of SNP used in the assay. Nevertheless, SNP antagonized the growth-promotion effect of gibberellin in a dose-dependent manner (Figure 1A and 1B). Treatment with GNSO showed a similar phenotype (Supplemental Figure 1A-1B). Consistent with these observations, the *gsnor1-3* mutant, which accumulates excessive amount of GSNO (Chen et al., 2009; Feechan et al., 2005; Lee et al., 2008), was nearly insensitive to gibberellin for the promotion effect on the elongation of hypocotyls (Figure 1C). Notably, gibberellin reduced the root growth of *gsnor1-3*, a phenotype opposite to that wild type (Figure 1D). These results suggest that NO antagonizes the gibberellin-promoted growth effect.

**Figure 1.**
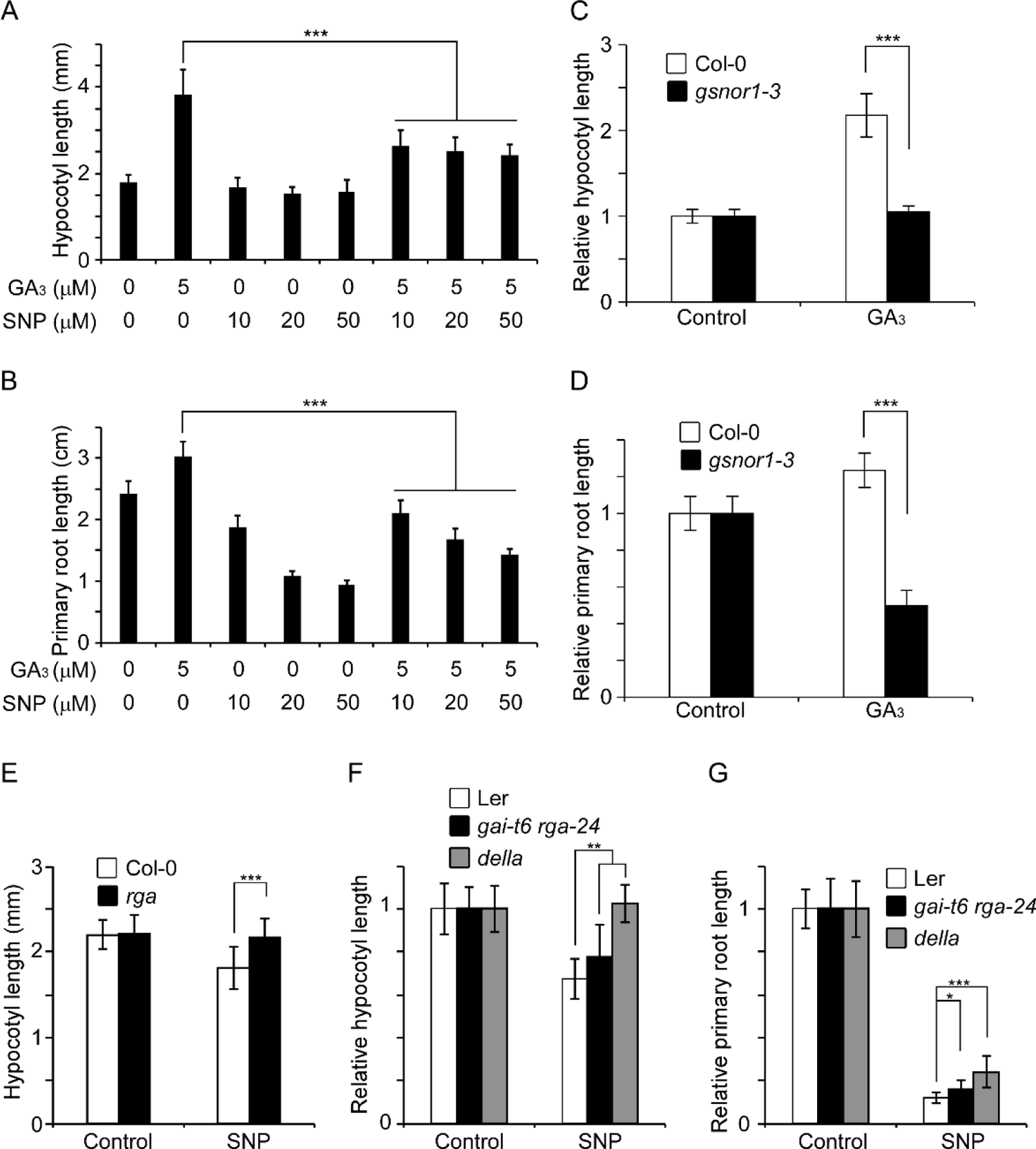
Nitric oxide antagonizes gibberellin-promoted root and hypocotyl elongation, see also Supplemental Figure S1 and S2. (A and B) Hypocotyl length (A) and primary root length (B) of 7-day-old Col-0 seedlings treated with the indicated concentrations of gibberellic acid (GA_3_) and sodium nitroprusside (SNP). (C and D) Hypocotyl length (C) and primary root length (D) of 7-day-old Col-0 and *gsnor1-3* seedlings treated with 5 μM GA_3_. (E) Hypocotyl length of 7-day-old Col-0 and *rga* seedlings treated with 20 μM SNP. (F and G) Hypocotyl length (F) and primary root length (G) of 7-day-old Col-0, *gai-t6 rga-24* and *della* (*gai-t6 rga-t2 rgl1-1 rgl2-1*) seedlings treated with 20 μM SNP. In each experiment, 30 seedlings were analyzed. *, **, and *** indicate *P* < 0.05, *P* < 0.01, and *P* < 0.001 (one-way ANOVA test), respectively.

We reasoned that NO may target DELLA repressor proteins to modulate gibberellin signaling. To test this possibility, we examined the response of mutants carrying various mutations in the *Arabidopsis DELLA* genes to NO. *Arabidopsis* has five *DELLA* genes, of which *RGA* and *GAI* are two major members (Dill et al., 2004; Schwechheimer and Willige, 2009; Xu et al., 2014). Among the analyzed mutants, *rga*, a T-DNA insertion mutant (SALK_089146), carries a null mutation (Supplemental Figure 2A-2C) and *della* is a quadruple mutant carrying null mutations in *RGA*, *GAI*, *RGL1*, and *RGL2* (Cheng et al., 2004). Under normal growth conditions, the *rga* mutant did not have detectable phenotype (Supplemental Figure 2D). However, the *rga* mutant was insensitive to the inhibitory effect of SNP on the elongation of hypocotyls (Figure 1E). Similarly, both the *gai-t6 rga-24* double mutant and *della* quadruple mutant were hyposensitive to SNP (Figure 1F and 1G). These results suggest that NO negatively regulates gibberellin signaling in a *DELLA*-dependent manner.

### Nitric oxide inhibits RGA-SLY1 interaction to stabilize RGA

Data presented above suggest that NO negatively regulates the gibberellin response via *DELLA* genes. We found that the transcription of *RGA* and key gibberellin biosynthesis genes was nearly unaltered when treated with *S*-nitrosoglutathione (GSNO) or SNP (Supplemental Figure 3A-B). Because the gibberellin-induced degradation of DELLA is a key step for the activation of gibberellin signaling, it is reasonable to assume that NO directly or indirectly regulates this class of repressor proteins. We then analyzed the regulation of NO on DELLA proteins. A *pRGA*::*GFP-RGA* transgenic line (Silverstone et al., 2001) was used to analyze the accumulation of RGA protein in response to NO. When treated with GSNO or SNO, the subcellular localization of GFP-RGA did not have detectable alterations (Supplemental Figure 3C). However, the accumulation of GFP-RGA protein was substantially increased by GSNO or SNP in a dose-dependent manner (Figure 2A and Supplemental Figure 3D). A time-course experiment revealed that the accumulation of GFP-RGA was progressively increased upon longer treatment with GSNO or SNP (Figure 2B and Supplemental Figure 3E). Consistent with these observations, the accumulation of RGA protein was significantly higher in NO over-accumulating mutant *gsnor1-3* and *nox1* (He et al., 2004) than that in wild type (Figure 2C), suggesting that NO positively regulates the stability of RGA. Remarkably, the gibberellin-induced degradation of RGA protein was nearly abolished by GSNO or SNP (Figure 2D and Supplemental Figure 3F), suggesting that NO inhibits gibberellin-promoted degradation of RGA protein.

**Figure 2.**
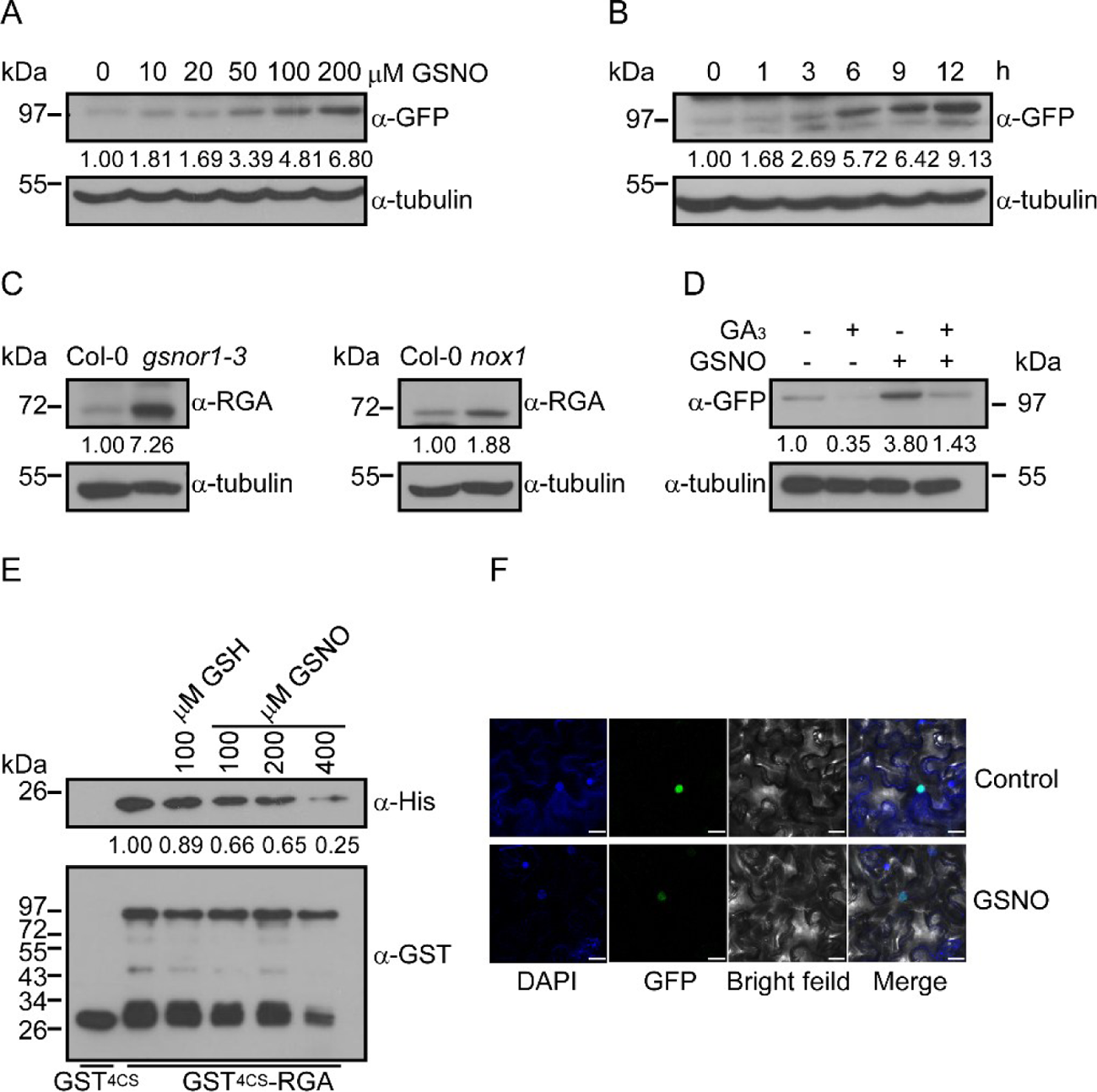
Nitric oxide inhibits gibberellin-promoted RGA degradation, see also Supplemental Figure S3, S4 and Table S1. (A and B) Immunoblotting analysis of GFP-RGA proteins in 7-day-old *pRGA*::*GFP-RGA* transgenic seedlings treated with indicated concentrations of GSNO for 6 hours (A) and 300 μM GSNO for the indicated times (B) by using an anti-GFP antibody. Immunoblotting with an anti-tubulin antibody is served as a loading control. Quantification of GFP-RGA is shown below the blot. (C) Immunoblotting analysis of RGA proteins in 7-day-old Col-0, *gsnor1-3*, and *nox1* seedlings by using an anti-RGA antibody. Quantification of RGA is shown below the blot. (D) Immunoblotting analysis of GFP-RGA proteins in 7-day-old *pRGA*::*GFP-RGA* transgenic seedlings treated with or without 300 μM GSNO and 0.5 μM GA_3_ for 6 hours using an anti-GFP antibody. Quantification of GFP-RGA is shown below the blot. (E) Analysis of the interaction of SLY1 and RGA1 recombinant proteins with a pull-down assay. GST^4CS^-RGA protein was treated with the indicated concentrations of GSNO or GSH prior to the incubation with His-SLY1. Quantification of the His-SLY1 level is shown below the blot. (F) Bimolecular fluorescence complementation (BiFC) analysis of co-localization of YNE-RGA1 and YCE-SLY1 fusion proteins in transiently expressed in tobacco leaves sprayed with 300 μM GSNO. Bar, 20 μm.

Upon binding to gibberellin, the activated GID1 receptor interacts with DELLA proteins to promote their association with the F-box protein SLY1, thereby facilitating the proteasomal degradation of the repressor proteins. To test if NO regulates the interaction between RGA-GID1 or RGA-SLY1, we performed the following experiments. We found that RGA recombinant protein physically interacted with SLY1 recombinant protein in a pull-down assay and the RGA-SLY1 interaction was reduced by GSNO in a dose-dependent manner (Figure 2E). A bimolecular fluorescence complementation (BiFC) assay also revealed that NO reduced the RGA-SLY1 interaction *in planta* (Figure 2F). Notably, the RGA-GID1 interaction is not regulated by NO (Supplemental Figure 4A and 4B). Taken together, these results suggest that NO positively regulates the stability of RGA by inhibiting its interaction with the F-box protein SLY1.

### *S*-nitrosylation of RGA at Cys-374 inhibits its proteasomal degradation

A major physiological role of NO is executed through protein *S*-nitrosylation. We then asked if RGA was posttranslationally modified by NO. We found that GSNO induced *S*-nitrosylation of RGA recombinant protein in an in vitro biotin-switch assay (Figure 3A). Similarly, GAI, RGL1, RGL2, and RGL3 recombinant proteins were also found being modified by *S*-nitrosylation (Supplemental Figure 5A-5D), suggesting that *S*-nitrosylation plays an important role in regulating DELLA proteins. Moreover, GFP-RGA protein was also found to be S-nitrosylated in planta (Figure 3B). Among 10 Cys residues in RGA, a mass spectrometric analysis of RGA recombinant protein identified Cys-249, Cys-374, Cys-506, and Cys-564 as *S*-nitrosylated residues (Figure 3C and Supplemental Table 1). In two replicates of mass spectrometry, four other Cys residues (Cys-228, Cys-286, Cys-299, and Cys-501) have also been covered, in which no modification was detected. However, Cys-129 and Cys-168 were not covered in mass spectrometry. We could not exclude the possibility that these two Cys residues are modified by *S*-nitrosylation. Because transgenic studies showed that mutations only in Cys-374, but not in Cys-249, Cys-506, and Cys-564, showed detectable effects under stress growth conditions (see below), we focused the analysis of Cys-374 hereafter and the functional studies of other *S*-nitrosylated Cys residues will be published elsewhere. Notably, Cys-374 is conserved in RGA and GAI, but not in other DELLA proteins, suggestive of possible functional divergence of these transcriptional repressors. The substitution of Cys-374 with Ser (RGA^C374S^) reduced the *S*-nitrosylation of the mutant protein in vitro and in planta (Figure 3D and 3E), indicating that Cys-374 is modified by *S*-nitrosylation.

**Figure 3.**
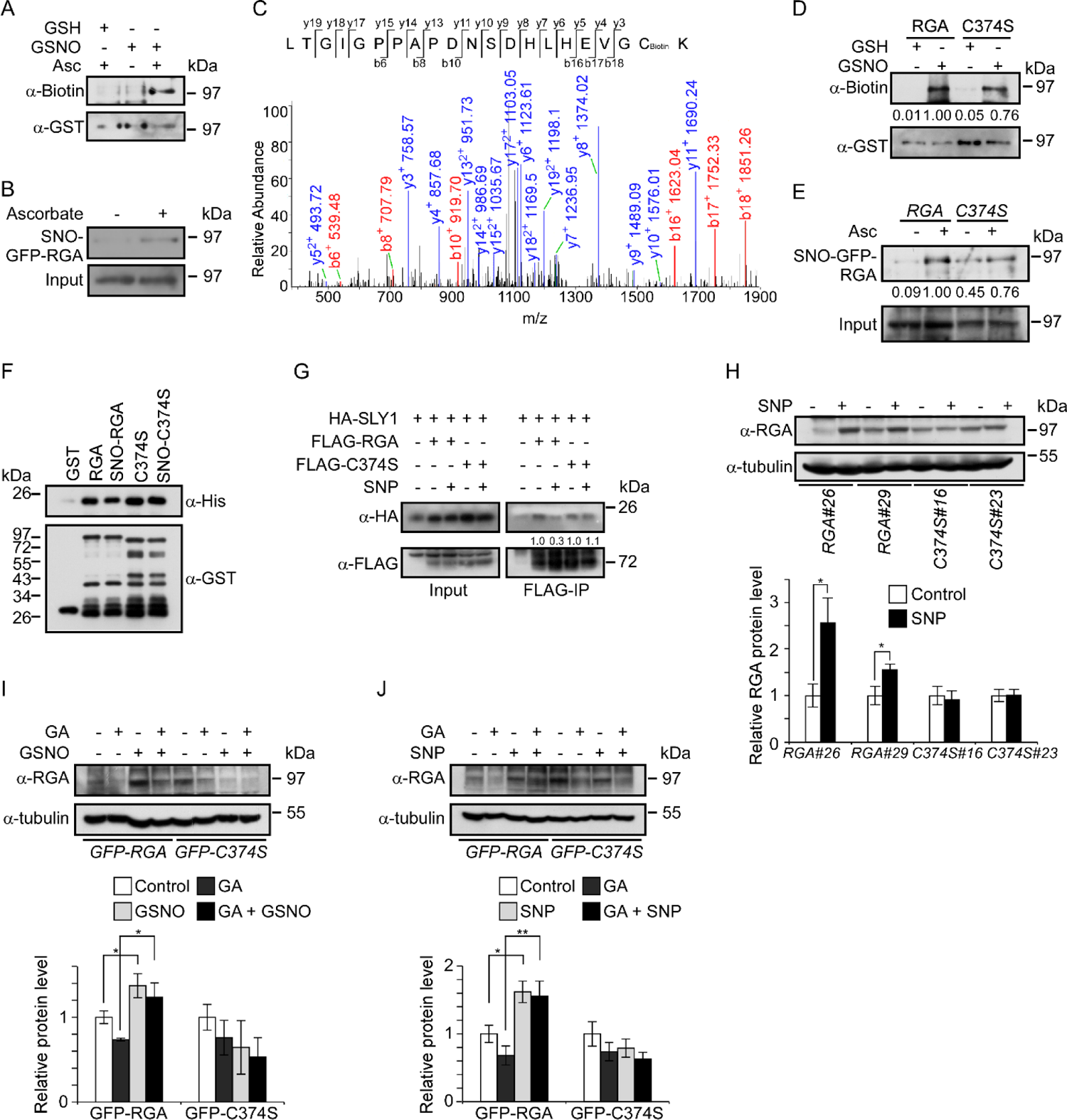
*S*-nitrosylation at Cys-374 modulates RGA stability, see also Supplemental Figure S4 and S5, Table S1 and Table S2. (A) Analysis of *S*-nitrosylated GST^4CS^-RGA recombinant protein treated with GSNO by an in vitro *S*-nitrosylation assay. Treatment with GSH and without sodium ascorbate (Asc) are served as negative controls. (B) Analysis of *S*-nitrosylated GFP-RGA protein in *pRGA*::*GFP-RGA* transgenic seedlings by an in vivo *S*-nitrosylation assay. The sample without Asc treatment is served as a negative control. (C) Liquid chromatography tandem-mass (LC-MS/MS) spectrum of trypsin-digested and biotin-charged RGA peptides. The b- and y-type product ions are indicated, which identified Cys-374 as an *S*-nitrosylated residue. (D) Analysis of *S*-nitrosylated GST^4CS^-RGA (RGA) and GST^4CS^-RGA^C374S^ (C374S) recombinant proteins treated with GSNO by an in vitro *S*-nitrosylation assay. Treatment with GSH is served as negative controls. Quantification of the *S*-nitrosylation level of GST^4CS^-RGA and GST^4CS^-RGA^C374S^ is shown below the blot. (E) Analysis of *S*-nitrosylated GFP-RGA and RGA^C374S^ (C374S) proteins in planta by an in vivo *S*-nitrosylation assay. The sample without Asc treatment is served as a negative control. Quantification of the *S*-nitrosylation level of RGA and RGA^C374S^ is shown below the blot. (F) Analysis of the interaction of RGA, RGA^C374S^ and SLY1 recombinant proteins with a pull-down assay. GST^4CS^-RGA and GST^4CS^-RGA^C374S^ was treated with 300 μM GSNO to generate *S*-nitrosylated proteins prior to the incubation with His-SLY1. (G) Analysis of the interaction of HA-SLY1, FLAG-RGA and FLAG-RGA^C374S^ (FLAG-C374S) proteins by a co-immunoprecipitation assay. The *HA-SLY1* and *FLAG-RGA* fusion genes under the control of the 35S promoter were transiently expressed in tobacco leaves that were incubated with or without 300 μM SNP for 1 hour. Protein extracts were used for Co-IP and analyzed by immunoblotting using anti-HA and -FLAG antibodies. Quantification of HA-SLY1 is shown below the blot. And protein level of HA-SLY1 that interacts with FLAG-RGA and FLAG-C374S without SNP treatment is set as 1.0, respectively. (H) Immunoblotting analysis of GFP-RGA proteins in *gai-t6 rga-24* transgenic seedlings of the indicated genotypes treated with 300 μM SNP for 6 hours using an anti-RGA antibody. Quantification of the GFP-RGA and GFP-RGA^C374S^ protein levels is shown below the blot. Protein levels of GFP-RGA and GFP-RGA^C374S^ without treatment are set as 1.0, respectively. (I) Immunoblotting analysis of GFP-RGA and GFP-RGA^C374S^ proteins in 7-day-old *pRGA*::*GFP-RGA*, *pRGA*::*GFP-RGA^C374S^* transgenic seedlings treated with or without 300 μM GSNO and 0.5 μM GA_3_ for 6 hours by using an anti-RGA antibody. Quantification of the GFP-RGA and GFP-RGA^C374S^ protein levels is shown below the blot. Protein levels of GFP-RGA and GFP-RGAC374S without treatment are set as 1.0, respectively. (J) Immunoblotting analysis of GFP-RGA and GFP-RGA^C374S^ proteins in 7-day-old *pRGA*::*GFP-RGA*, *pRGA*::*GFP-RGA^C374S^* transgenic seedlings treated with or without 300 μM SNP, and 0.5 μM GA_3_ for 6 hours by using an anti-RGA antibody. Quantification of the GFP-RGA and GFP-RGA^C374S^ protein levels is shown below the blot. Protein levels of GFP-RGA and GFP-RGA^C374S^ without treatment are set as 1.0, respectively. Data presented in (H)-(J) are means of three independent experiments with S.D. * and ** indicate *P* < 0.05 and *P* < 0.01, respectively (One-way ANOVA test).

Because the interaction of RGA with SLY is negatively regulated by NO, we reasoned that the RGA-SLY interaction might be regulated by *S*-nitrosylation. While the RGA-SLY1 interaction was reduced by GSNO, this negative effect was abolished by a *RGA^C374S^* mutation in a pull-down assay (Figure 3F). Moreover, the interaction of SLY1 and RGA was detected by a co-immunoprecipitation (Co-IP) assay when transiently expressed in tobacco (*Nicotiana tabacum*) leaves harboring *HA-SLY1* and *FLAG-RGA* or *FLAG-RGA^C374S^* constructs and the interaction was inhibited by SNP. However, SNP exerts NO inhibitory effect on RGA^C374S^-SLY1 interaction (Fig 3G). Collectively, these results suggest that *S*-nitrosylation at Cys-374 negatively regulates the RGA-SLY1 interaction. Consistent with this observation, the SNP-induced accumulation of GFP-RGA protein was abolished by the *RGA^C374S^* mutation (Figure 3H). Moreover, the gibberellin-induced degradation of RGA was inhibited by SNP and GSNO in RGA, but not in RGA^C374S^ mutant proteins (Figure 3I and 3J), suggesting that Cys-374 confers the responsiveness of RGA to NO. These results suggest that *S*-nitrosylation of RGA at Cys-374 inhibits its interaction with the F-box protein SLY1, thereby preventing its proteasomal degradation.

### *S-*nitrosylation of RGA at Cys-374 coordinates plant growth and stress responses

Given the importance of *S*-nitrosylation in regulating the stability of RGA, we next explored its physiological significance in modulating growth and stress responses. To this end, a *pRGA*::*GFP-RGA^C374S^* transgene and its control *pRGA*::*GFP*-*RGA* were introduced into the *gai-t6 rga-24* double mutant by genetic transformation. The *gai-t6 rga-24* double mutant showed elongated hypocotyls and primary roots under normal growth conditions and this phenotype was fully rescued by both *pRGA*::*GFP*-*RGA* and *pRGA*::*GFP*-*RGA^C374S^* transgenes (Figure 4A-4C and Supplemental Figure 1B). Moreover, these two transgenes also fully rescued the sensitivity of the *gai-t6 rga-24* double mutant to gibberellin (Figure 4A-4C and Supplemental Figure 1A).

**Figure 4.**
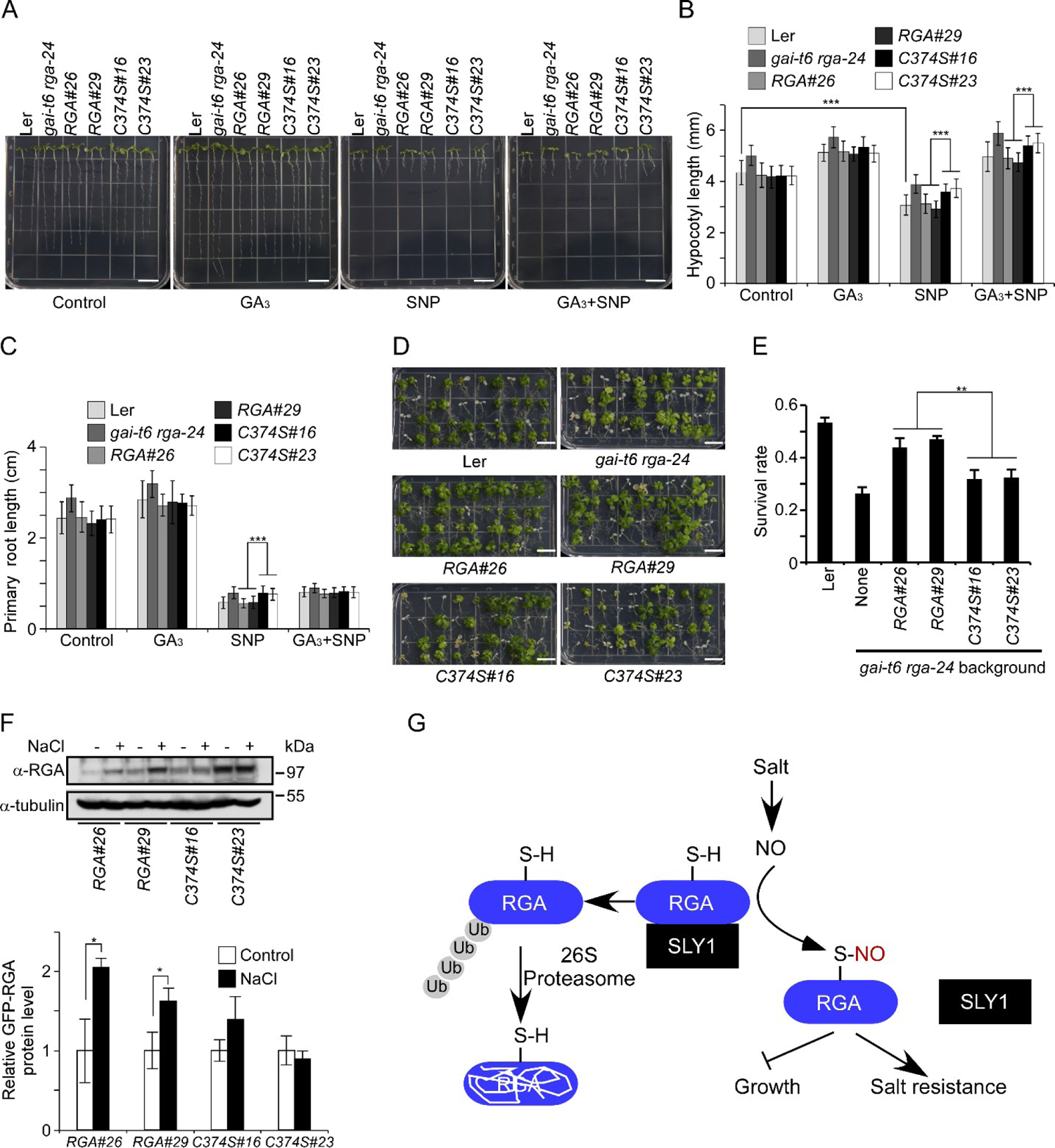
*S*-nitrosylation of RGA balances plant growth and salinity tolerance, see also Figure S4 and Supplemental Table S2. (A) Ten-day-old seedlings of the indicated genotypes treated with 50 μM SNP or 5 μM GA_3_. Bar, 1 cm. (B) and (C) Analysis of hypocotyl length (B) and primary root length (C) of transgenic seedlings of the indicated genotypes treated with 50 μM SNP or 5 μM GA_3_ for 10 days. (D) Five-day-old seedlings of the indicated genotypes on 1/2MS medium were transferred to the medium containing 125 mM NaCl. Photos were taken 2 weeks post the transfer. Bar, 1 cm. (E) Analysis of the survival rate of transgenic seedlings of the indicated genotypes shown in (D). (F) Immunoblotting analysis of GFP-RGA proteins in *gai-t6 rga-24* transgenic seedlings of the indicated genotypes treated with 150 mM NaCl for 6 hours using an anti-RGA antibody. Quantification of the GFP-RGA and GFP-RGA^C374S^ protein levels is shown below the blot. (G) A proposed model illustrating the function of *S*-nitrosylation of RGA. Under normal growth conditions, SLY1 interacts with RGA, which leads to the polyubiquitination and degradation of RGA via the 26S proteasome pathway. High salt induces the NO burst, which subsequently induces the *S*-nitrosylation of RGA. The *S*-nitrosylation inhibits the RGA-SLY1 interaction and enhances the stability of RGA. Accumulating RGA inhibits plant growth and enhance the salinity tolerance. ≥ 30 seedlings are analyzed for each sample in (B) and (C). Data presented in (E) and (F) are means of three independent experiments with S.D. * and ** indicate *P* < 0.05 and *P* < 0.01, respectively (One-way ANOVA test).

However, the response of *gai-t6 rga-24* to SNP and GSNO, regardless of the presence or the absence of gibberellin, was only rescued by *pRGA*::*GFP*-*RGA*, but not by *pRGA*::*GFP*-*RGA^C374S^* (Figure 4A-4C, Supplemental Figure 1), consistent with the observation that the accumulation of RGA, but not RGA^C374S^, was sensitive to SNP and GSNO (see Figure 3H-3J). These results suggest that *S*-nitrosylation of RGA at Cys-374 plays an important role in regulating gibberellin signaling.

While NO is a key regulator of stress responses (Astier et al., 2011; Feng et al., 2019; Yu et al., 2014), gibberellin signaling is also implied to play a role in salt tolerance, evidenced by the observation that the *gai-t6 rga-24* mutant is hypersensitive to NaCl (Achard et al., 2006). We then asked if *S*-nitrosylation of RGA is involved in regulating stress responses. We found that the hypersensitivity of *gai-t6 rga-24* to NaCl was restored by *pRGA*::*GFP*-*RGA*, but not by *pRGA*::*GFP*-*RGA^C374S^* (Figure 4D-4E). This phenotype was correlated to the accumulation of RGA and RGA^C374S^ proteins in response to NaCl (Figure 4F), in a manner similar to that of SNP (see Figure 3H), suggesting that *S*-nitrosylation of RGA at Cys-374 is essential for its responsiveness to a stress signal. Taken together, these results suggest that *S*-nitrosylation of RGA modulates gibberellin signaling and stress tolerance to coordinate plant growth in response to variable environmental conditions (Figure 4G).

## Discussion

In this study, we find that NO negatively regulates gibberellin signaling by stabilizing the RGA repressor via *S*-nitrosylation, which retards growth but positively modulates stress tolerance, thus uncovering a unique mechanism balancing the growth and survival of plants (Figure 4G). While gibberellin is a key regulator promoting plant growth in most, if not all, developmental stages, the burst of NO is generally believed as a hallmark at the onset of stress responses. When challenged by environmental stress, plants usually respond by the inhibition of growth and the activation of stress responses to cope with the detrimental growth conditions. It has been recognized that NO boosts stress tolerance via *S*-nitrosylation of key components of stress responses (Albertos et al., 2015; Hu et al., 2017; Wang et al., 2015; Yang et al., 2015). Also as a protective mechanism, stresses promote the accumulation of DELLA proteins, mediated by decreasing the biosynthesis of gibberellins (Achard et al., 2006; Wang et al., 2020b) or repressing the transcription of *SLY1*, encoding an F-box-containing E3 ligase directly mediating the proteasomal degradation of DELLAs (Lozano-Juste and Leon, 2011), which causes growth inhibition. However, while the inhibitory role of NO on plant growth has been noticed, the underpinning mechanisms remains largely unknown. The finding that *S*-nitrosylation of RGA, a major member of DELLA repressor proteins, inhibits its interaction with SLY1 and consequently prevent its proteasomal degradation, reveals a unique regulatory mechanism that confers plants a more rapid and efficient response when sensing adverse growth conditions. Moreover, we also find that *S*-nitrosylation of RGA at Cys-347 is essential for its regulatory role in salt stress responses. Together, the NO-mediated *S*-nitrosylation of RGA inhibits growth whereas enhances salt stress tolerance, representing a unique mechanism that balances the growth and survival of plants when challenged by detrimental growth conditions.

While an interplay between gibberellin and NO signaling coordinates plant growth and stress responses as revealed in this study, an analogous regulatory scheme has also been appreciated between the cytokinin and NO pathways. The *S*-nitrosylation of AHP1, a key regulator of cytokinin responses, causes a reduction of its phosphorylation, thereby negatively regulating signaling of this growth-promotion phytohormone (Feng et al., 2013). Therefore, *S*-nitrosylation may represent an important mechanism that integrates an NO signal into signaling of growth-promotion phytohormones and eventually retards growth in response to environmental stresses. It has been noticed that the protein *S*-nitrosylation level is tightly regulated by the intracellular NO concentrations (Benhar et al., 2009; Feng et al., 2019; Hess et al., 2005; Hu et al., 2015), which is induced by various stimuli in fluctuating environments (Wang et al., 2010; Zhao et al., 2009; Zhou et al., 2016). Thus, *S*-nitrosylation of DELLA proteins permits a rapid response to diverse environmental alterations. When growth conditions become favorable, the intracellular NO level is returned to a physiologically normal level, which may trigger a reverse denitrosylation reaction (Benhar et al., 2009; Kneeshaw et al., 2014; Tada et al., 2008). It is reasonable to speculate that the denitrosylation of RGA resets the transcriptional repressor under the control of gibberellin-promoted proteasomal degradation, thereby relieving from the growth retardation. Therefore, RGA acts a signaling molecule to sense intracellular NO level to coordinate plant growth and stress tolerance in response to dynamically altered environment.

In addition to Cys-374, several other Cys residues, including Cys-249, Cys-506, and Cys-564, of RGA are also identified as the *S*-nitrosylated sites in mass spectrometric analysis. While *S*-nitrosylation of these Cys residues remains functionally unclear, it is well known that gibberellins regulate multiple biological processes, largely dependent on the interactions between DELLAs and transcription factors of other signaling pathways. For instance, GAI and RGA interact physically with PIF3 and PIF4, two bHLH transcription factors, to regulates the plant growth (de Lucas et al., 2008; Feng et al., 2008). As NO is invovled in regulating diverse biological processes, it is of great interest to investigate whether *S*-nitrosylation at Cys-249, Cys-506, or Cys-564 affects interaction between RGA and its interacting proteins.

Finally, DELLA proteins are regulated by multiple forms of posttranslational modifications, including phosphorylation, SUMOylation, *O*-GlcNAcylation, and *O*-fucosylation (Conti et al., 2014; Dai and Xue, 2010; Zentella et al., 2016). Investigation of possible interactions of these posttranslational modifications, including *S*-nitrosylation, will be of great interests toward the understanding how plants balance growth and stress tolerance in response to environmental alterations.

## SUPPLEMENTAL INFORMATION

Supplemental information includes 5 figures and 2 tables and can be found with this article online.

## ACKNOWLEDGEMENTS

We thank the *Arabidopsis* Biological Resource Center (ABRC) and Xiangdong Fu for seeds. This work was supported by grants from the National Natural Science Foundation of China (31830017 and 31521001), Chinese Academy of Sciences (XDB27030207), and State Key Laboratory of Plant Genomics (SKLPG2020-22).

## AUTHOR CONTRIBUTIONS

L.C. performed the majority of the experiment, assisted by S.S. J.Z., L.C., J.-M.Z., and JL designed the experiments and analyzed the data. J.Z. wrote the manuscript, assisted by L.C. All authors discussed the results and commented on the manuscript.

## STAR*METHODS

### KEY RESOURCES TABLE

| REAGENT or RESOURCE | SOURCE | IDENTIFIER |
| --- | --- | --- |
| Antibodies |  |  |
| HRP-linked anti-biotin | Cell Signaling Technology | Cat#7075 |
| Mouse monoclonal anti-GFP | Abmart | Cat# M20004L |
| Mouse monoclonal anti-GST | Abgent | Cat # AM1011a |
| Mouse monoclonal anti-His | CMCTAG | Cat # AT0025 |
| Mouse monoclonal anti-tubulin | Sigma-Aldrich | Cat# T5168 |
| Rabbit polyclonal anti-HA | Abgent | Cat# AP1012a |
| Mouse monoclonal anti-FLAG<sup>®</sup> M2-peroxidase (HRP) antibody | Sigma-Aldrich | Cat# A8592 |
| Mouse polyclonal anti-RGA | This paper | N/A |
| Bacterial and Virus Strains |  |  |
| *E. coli* DH5 | TransGen Biotech | Cat# CD201 |
| *E. coli* BL21 | TransGen Biotech | Cat# CD901 |
| *Agrobacterium tumefaciens* GV3101 | Biomed | Cat# BC304 |
| Chemicals, Peptides, and Recombinant Proteins |  |  |
| His-SLY1 | This paper | N/A |
| His-GID1a | This paper | N/A |
| His-GID1c | This paper | N/A |
| His-RGA Nter | This paper | N/A |
| GST<sup>4CS</sup> | (Feng et al., 2013) | N/A |
| GST<sup>4CS</sup>-RGA | This paper | N/A |
| GST<sup>4CS</sup>-RGA<sup>C374S</sup> | This paper | N/A |
| GST<sup>4CS</sup>-GAI | This paper | N/A |
| GST<sup>4CS</sup>-RGL1 | This paper | N/A |
| GST<sup>4CS</sup>-RGL2 | This paper | N/A |
| GST<sup>4CS</sup>-RGL3 | This paper | N/A |
| S-nitrosoglutathione (GSNO) | Sigma-Aldrich | Cat# N4148 |
| Sodium nitroprusside (SNP) | Sigma-Aldrich | Cat# 71778 |
| Biotin-HPDP | Thermo Scientific | Cat# 21341 |
| Biotin-maleimide | Sigma-Aldrich | Cat# B1267 |
| Glutathione | Sigma-Aldrich | Cat# G4251 |
| Methyl methanethiosulfonate (MMTS) | Thermo Scientific | Cat# 23011 |
| Neocuproine | Sigma-Aldrich | Cat# N1501 |
| Sodium ascorbate | Sigma-Aldrich | Cat# A7631 |
| Gibberellic acid (GA<sub>3</sub>) | TCI | Cat# G0029 |
| Paclobutrazol | Sigma-Aldrich | Cat# 46046 |
| Protease Inhibitor Cocktail | Sigma-Aldrich | Cat# P9599 |
| Trypsin | Promega | Cat# V5280 |
| Critical Commercial Assays |  |  |
| Ni-NTA resin | QIAGEN | Cat# 30210 |
| Glutathione Sepharose | GE | Cat# 17-0756-01 |
| High Capacity Neutravidin Agarose Resin | Thermo Scientific | Cat# 29202 |
| Anti-FLAG® M2 Affinity Gel | Sigma-Aldrich | Cat# A2220 |
| Zeba Spin Desalting Columns | Thermo Scientific | Cat# 89883 |
| Experimental Models: Organisms/Strains |  |  |
| *Arabidopsis: gsnor1-3* | (Feechan et al., 2005) | GABI_315D11 |
| *Arabidopsis: nox1* | (He et al., 2004) | N/A |
| *Arabidopsis: rga* | This paper | SALK_089146 |
| *Arabidopsis: gai-t6 rga-24* | (King et al., 2001) | N/A |
| *Arabidopsis: gai-t6 rga-t2 rgl1-1 rgl2-1* | (Achard et al., 2006) | N/A |
| *Arabidopsis: pRGA:: GFP-RGA* | (Silverstone et al., 2001) | N/A |
| *Arabidopsis: pRGA:: GFP-RGA (pER8)* | This paper | N/A |
| *Arabidopsis: pRGA:: GFP-RGA<sup>C374S</sup>* | This paper | N/A |
| Oligonucleotides |  |  |
| Primers for cloning, PCR and genotyping | This paper (see Table S1) | N/A |
| Recombinant DNA |  |  |
| pET28a-SLY1 | This paper | N/A |
| pET28a-GID1a | This paper | N/A |
| pET28a-GID1c | This paper | N/A |
| pET28a-RGA Nter | This paper | N/A |
| pGST<sup>4CS</sup> | (Feng et al., 2013) | N/A |
| pGST<sup>4CS</sup>-RGA | This paper | N/A |
| pGST<sup>4CS</sup>-RGA<sup>C374S</sup> | This paper | N/A |
| pGST<sup>4CS</sup>-GAI | This paper | N/A |
| pGST<sup>4CS</sup>-RGL1 | This paper | N/A |
| pGST<sup>4CS</sup>-RGL2 | This paper | N/A |
| pGST<sup>4CS</sup>-RGL3 | This paper | N/A |
| pWM101-FLAG-RGA | This paper | N/A |
| pWM101-FLAG-RGA<sup>C374S</sup> | This paper | N/A |
| pWM101-HA-SLY1 | This paper | N/A |
| pER8-pRGA:: GFP-RGA | This paper | N/A |
| pER8-pRGA:: GFP-RGA<sup>C374S</sup> | This paper | N/A |
| Software and Algorithms |  |  |
| ImageJ | NIH | http://imagej.nih.gov/ij/ |

### LEAD CONTACT AND MATERIALS AVAILABILITY

Further information and requests for resources and reagents should be directed to and will be fulfilled by the Lead Contact, Jianru Zuo

### EXPERIMENTAL MODEL AND SUBJECT DETAILS

Columbia-0 (Col-0) and Landsberg *erecta* (L*er*) accessions of *Arabidopsis* was used in this study. The *gsnor1-3* mutant seeds (Feechan et al., 2005) were provided by Gary Loake. The *nox1* mutant seeds (He et al., 2004) were provided by Yikun He. The *gai-t6 rga-24*, *gai-t6 rga-t2 rgl1-1 rgl2-1*, and *pRGA*::*GFP-RGA* transgenic line (L*er* background) (Achard et al., 2006; King et al., 2001; Silverstone et al., 2001) were provided by Xiangdong Fu. The *rga* (SALK_089146) mutant was obtained from ABRC. Generation of transgenic *Arabidopsis* plants was carried out by *Agrobacterium*-mediated transformation (Bechtold and Pelletier, 1998). The *pRGA*:: *GFP-RGA* and *pRGA*::*GFP-RGA^C374S^* were introduced into *gai-t6 rga-24* plants. T2 or subseqent generations of transgenics that are homozygous for a single insertion were used for all studies. At least two independent transgenic lines are analyzed. Unless specified otherwise, no apparent phenotype was observed in these transgenic plants under normal growth conditions.

Seeds were sterilized and sown on 1/2 MS medium agar plates with 1% sucrose. The seeds were imbibed at 4°C for 2 days, and then cultured at 22°C under 16/8 h light/dark.

## METHODS DETAILS

### Plasmid construction

The coding sequence of *RGA* was inserted into the *BamH*I/*Sal*I sites of pGST^4CS^, a modified pGEX4T1 vector, to generate pGST^4CS^-RGA. The pGST^4CS^-GAI, pGST^4CS^-RGL1, pGST^4CS^-RGL2, and pGST^4CS^-RGL3 constructed were generated in a similar way. The coding sequences of *SLY1* and *GID1a* were inserted into the *BamH*I/*Hind*III sites and *BamH*I/*Sal*I sites, respectively, to produce pET28a-SLY1 and pET28a-GID1a. The pET28a-GID1c and pET28a-RGA Nter vectors were generated in a similar way.

Putative promoter sequences and the coding regions of *RGA* were separately amplified and appropriate restriction sites were introduced during PCR. The promoter fragment was digested with *Xho*I/*Nco*I, and cloned into pSK-GFP with the same sites to generate pSK-pRGA::GFP. Fragments of *pRGA*::*GFP* and *RGA* coding regions were digested with *Xho*I/*Pst*I and *Pst*I/*Spe*I, respectively, and then cloned into the *Xho*I/*Spe*I sites of a pER8 binary vector (Zuo et al., 2000) to generate pER8-pRGA::GFP-RGA.

The coding sequences of *RGA* and *SLY1* were PCR-amplified and in-frame fused to an FLAG or HA tag to yield pSK-FLAG-RGA and pSK-HA-SLY1, respectively. *FLAG-RGA* and *HA-SLY1* were PCR-amplified and ligated to pMW101 at *Kpn*I/*Sal*I and *Kpn*/*Pst*I under the control of a 35S promoter, respectively, to generate pWM101-FLAG-RGA and pWM101-HA-SLY1.

The BiFC expression vectors pCAMBIA1300-YNE-RGA and pCAMBIA1300-YCE-SLY1 were constructed by a approach described previously (Chen et al., 2020).

Site-directed mutagenesis was performed using the Easy Mutagenesis System (TransGen Biotech, Beijing) according the manufacturer’s instructions. Primers used for the mutagenesis are listed in Table S1.

All constructs were verified by extensive restriction digestion and DNA sequencing analysis. All PCR-related primers used in this study are listed in Table S1.

### Expression and purification of recombinant proteins

The pET28a-SLY1, pET28a-GID1a, pET28a-GID1c, pET28a-RGA Nter, pGST^4CS^-RGA, pGST^4CS^-RGA ^C374S^, pGST^4CS^-GAI, pGST^4CS^-RGL1, pGST^4CS^-RGL2, and pGST^4CS^-RGL3 expression vectors were transformed into *Escherichia coli* strain BL21 (DE3). Expression and purification of the recombinant proteins were carried out following the manufacturer’s instructions.

### Generation of antibodies and immunoblotting

Anti-tubulin (Sigma-Aldrich, Cat #T5168), anti-GST (Abgent, Cat #AM1011a), anti-His (CMCTAG, Cat #AT0025), anti-GFP (Abmart, Cat#M20004L), anti-HA (Abgent, Cat #AP1012a) and anti-FLAG (Sigma-Aldrich, Cat #A8592) antibodies were obtained from commercial sources. RGA-specific antibodies were generated by immunizing mice with *Escherichia coli*-expressed N-terminal of RGA (His-RGA Nter). Total protein was extracted and no cross-reaction was observed in the *rga* mutant compared with Col-0 (Supplemental Figure 2C). Immunoblotting was carried out as previously described (Chen et al., 2009). Quantification of the immunoblot was performed using NIH ImageJ (version 1.44p; http://imagej.nih.gov/ij/).

### Bimolecular fluorescence complementation analysis

Bimolecular fluorescence complementation (BiFC) assays were performed as described (Chen et al., 2020). The *Nicotiana benthamiana* leaves were injected with agrobacteria cultures containing expression vectors and cultured for additional two days. 300 μM GSNO was sprayed on the surface of the tobacco leaves for 2 hours. The leaves were then excised and observed under a confocal microscope.

### In vitro pull down

GST^4CS^-tagged RGA or RGA^C374S^ recombinant proteins were incubated with indicated concentrations of GSH or GSNO in HEN buffer (250mM Hepes, pH 7.7, 1mM EDTA, 1 mM neocuproine) for 30 min. Free GSNO or GSH was removed using Zeba Spin Desalting Columns (Thermo, Cat #: 89883). 1 μg GST^4CS^ and 2 μg GST^4CS^-tagged RGA or RGA^C374S^ recombinant proteins immobilized on Glutathione Sepharose. Immobilized beads were incubated with 1 μg His-tagged recombinant proteins in PBS-140N buffer (137 mM NaCl, 2.7 mM KCl, 10 mM Na_2_PO_4_, 2 mM KH_2_PO_4_, pH 7.4, 0.5% IGEPAL CA-630) for 1 hr at 4°C. For His-GID1a and His-GID1c, 10 μM GA_3_ was added. The supernatant was removed after centrifugation at 800 rpm, and the beads were washed six times with precooled PBS-140N buffer. The resin-retained proteins were analyzed by western blot analysis using anti-His or anti-GST antibodies as indicated.

### Co-immunoprecipitation

Co-immunoprecipitation experiments were performed as previously described (Ren et al., 2013) with modifications. pWM101-FLAG-RGA, pWM101-FLAG-RGA^C374S^, and pWM101-HA-SLY1 constructs were transiently expressed in tobacco leaves by agrobacterium-mediated infiltration (strain GV3101). After cultured for additional three days, tobacco leaves were incubated with 10 μM paclobutrazol for 3 hours and treated with 300 μM SNP for another 1 hour. Tobacco leaves were then ground in liquid nitrogen and extracted in IP buffer (50 mM Tris-HCl, pH 7.4, 150 mM NaCl, 5% glycerol, 0.1% IGEPAL CA-630) supplemented with Protease Inhibitor Cocktail. Samples were centrifuged at 13,000 rpm for 20 min at 4°C and the supernatant was collected. Proteins were incubated with anti-FLAG^®^ M2 affinity gel (Sigma, Cat # A2220) for 2 hours at 4°C. The gel were washed 6 times with IP buffer and proteins were then eluted and analyzed by immunoblotting.

### In vitro *S*-nitrosylation assay

Analysis of *in vitro S*-nitrosylation was performed essentially as described (Chen et al., 2020). Approximately 10 μg of GST^4CS^-tagged RGA or RGA^C374S^ recombinant proteins were incubated with GSNO or GSH at a final concentration of 200 μM in the dark for 30 min. Protein was precipitated by adding three volumes of cold acetone. The pellet was washed three times with 70% acetone and resuspended in 200 μL blocking buffer 1 (250 mM Hepes, pH 7.7, 4 mM EDTA, 1 mM neocuproine, 2.5% SDS and 200 mM *S*-methylmethane thiosulfonate). After incubation at 50°C for 40 min, protein was precipitated by adding three volumes of acetone and washed with 70% acetone. The pellet is dissolved in 80 μL HENS buffer (250 mM Hepes, pH 7.7, 4 mM EDTA, 1 mM neocuproine, 1% SDS), followed by addition of 10 μL 500 mM sodium ascorbate and 10 μL of 4 mM biotin-HPDP. The reaction was run for 1 hr at room temperature. Samples were separated by SDS-PAGE and analyzed by immunoblotting using an anti-biotin antibody (Cell Signaling Technology, Cat#7075).

### In vivo *S*-nitrosylation assay

Analysis of *in vivo S*-nitrosylation was performed as described (Feng et al., 2013) with minor modifications. In brief, two-week-old seedlings were ground in liquid nitrogen and extracted HEN/RIPA buffer (250 mM Hepes, pH 7.7, 1mM EDTA, 0.1 mM neocuproine, 1% Triton X-100, protease inhibitor cocktail, 0.1% SDS and 1% sodium deoxycholate). 300 μg protein was incubated with blocking buffer at 50°C for 40 min. Protein was precipitated with cold acetone.

The pellet was washed three times with 70% acetone and resuspended in 240 μL of HENS buffer followed by addition of 30 μL of 500 mM sodium ascorbate and 30 μL of 4 mM biotin-HPDP. The reaction was run for 1 hr at room temperature. Protein was precipitated with cold acetone, washed three times with 70% acetone and resuspended in 300 μL HENS buffer. After being neutralized with 900 μL of neutralization buffer (25 mM HEPES, pH 7.7, 100 mM NaCl, 1 mM EDTA, and 0.5% Triton X-100), the sample was mixed with 40 μL of streptavidin beads (Thermo Scientific, Cat #29202) and incubated at 4°C overnight. The beads were washed six times with washing buffer (25 mM HEPES, pH 7.7, 600 mM NaCl, 1 mM EDTA, and 0.5% Triton X-100). The proteins were then eluted and analyzed by immunoblotting.

### Mass spectrometric analysis of *S*-nitrosylation residues

Mass spectrometric identification of *S*-nitrosylated cysteine residues was carried out as described (Chen et al., 2020). Approximately 30 μg GST^4CS^-RGA recombinant proteins were labeled with biotin-maleimide (Sigma-Aldrich, Cat#B1267). The biotinylated protein was digested with Trypsin (Promega, Cat#V5280) in gel. The Trypsin-digested sample was analyzed by LC-MS/MS using a Thermo Fisher Finnigan linear ion trap quadrupole mass spectrometer in line with a Thermo Fisher Finnigan Surveyor MS Pump Plus HPLC system. The raw data was searched against the GST^4CS^-RGA protein sequence using pFIND searching software. Cysteine biotinylation (451.200 Da), cysteine carbamidomethylation (57 Da), and methionine oxidation (15.995 Da) were included in the search as the variable modifications.

## QUANTIFICATION AND STATISTICAL ANALYSIS

For quantification analyses, the mean and SD were calculated and compared to control and significance (*P* value) was determined using the two-tailed Student’s *t*-test or one-way ANOVA test (specified in Figure legends). All experiments were repeated at least 3 times, and representative results are shown.

---

**Figure S1.**
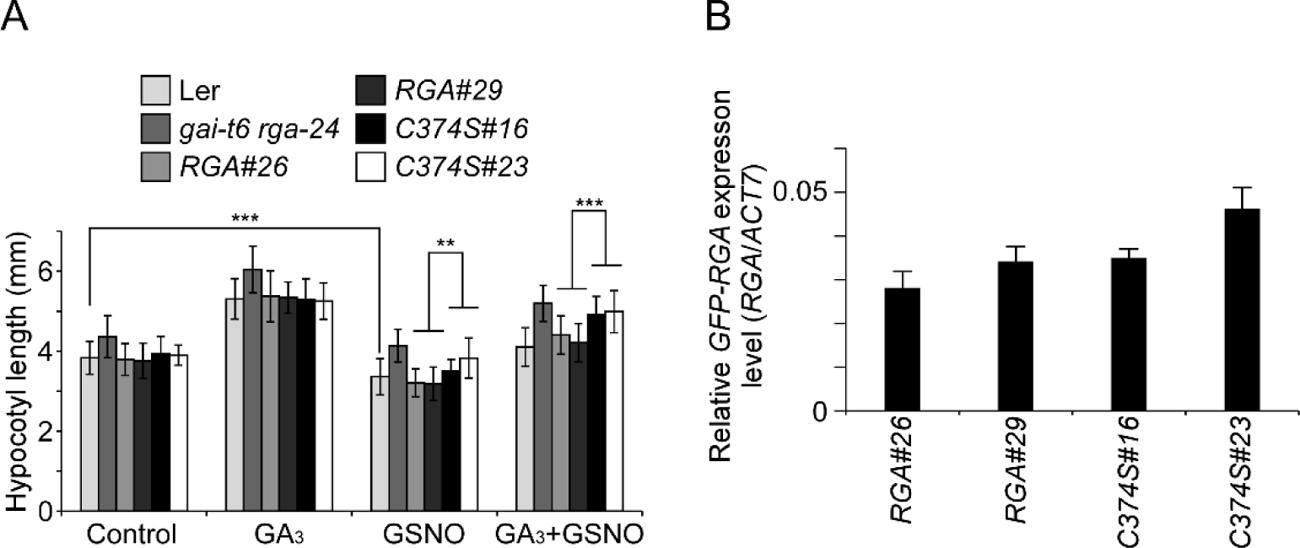
*S*-nitrosylation of RGA modulates plant growth, related to Figure 1 and 4. (A) Analysis of hypocotyl length of transgenic seedlings of the indicated genotypes treated with 300 μM GSNO or 5 μM GA_3_ for 10 days. Thirty seedlings were analyzed. *, **, and *** indicate *P* < 0.05, *P* < 0.01, and *P* < 0.001 (one-way ANOVA test), respectively. (B) Analysis of the expression of *GFP-RGA* by qRT-PCR in 7-day-old transgenic seedlings.

**Figure S2.**
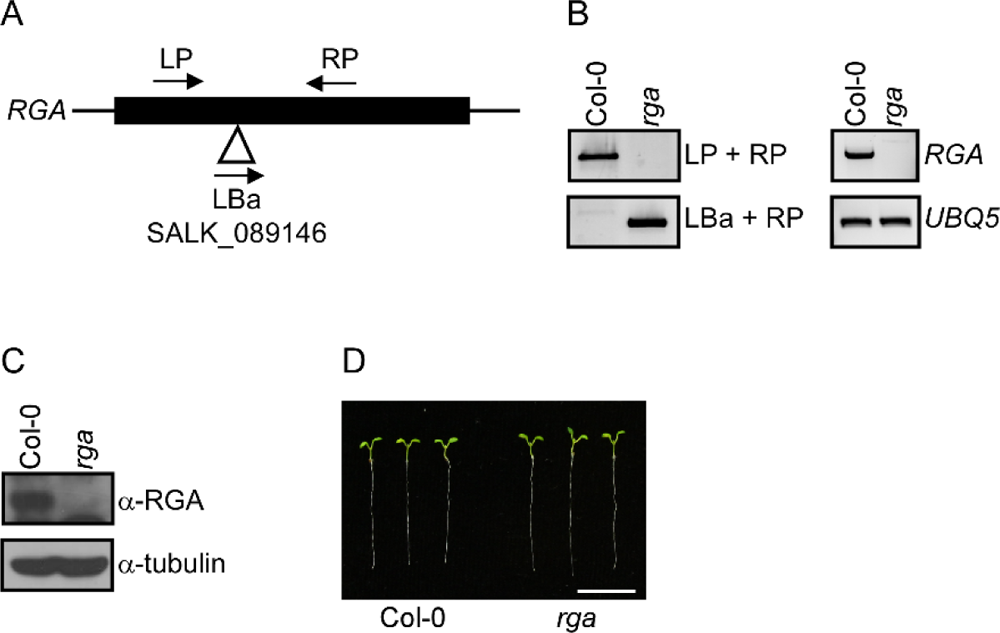
Characterization of *rga* mutant, related to Figure 1. (A) The gene structure is indicated with exons represented by boxes and UTRs represented by lines. The position of T-DNA is indicated with open triangle. (B) Genotyping of *rga* mutant. (C) Analysis of *RGA* expression by RT-PCR. *RGA* expression in Col-0 and the *rga* mutant is shown. *UBQ5* was used as loading control. (D) Analysis of RGA protein level in Col-0 and the *rga* mutant. Total protein was extracted from 7-day-old seedlings and probed with anti-RGA and anti-tubulin antibodies. (E) Seven-day-old seedlings of Col-0 and *rga* mutant. Bar, 1 cm.

**Figure S3.**
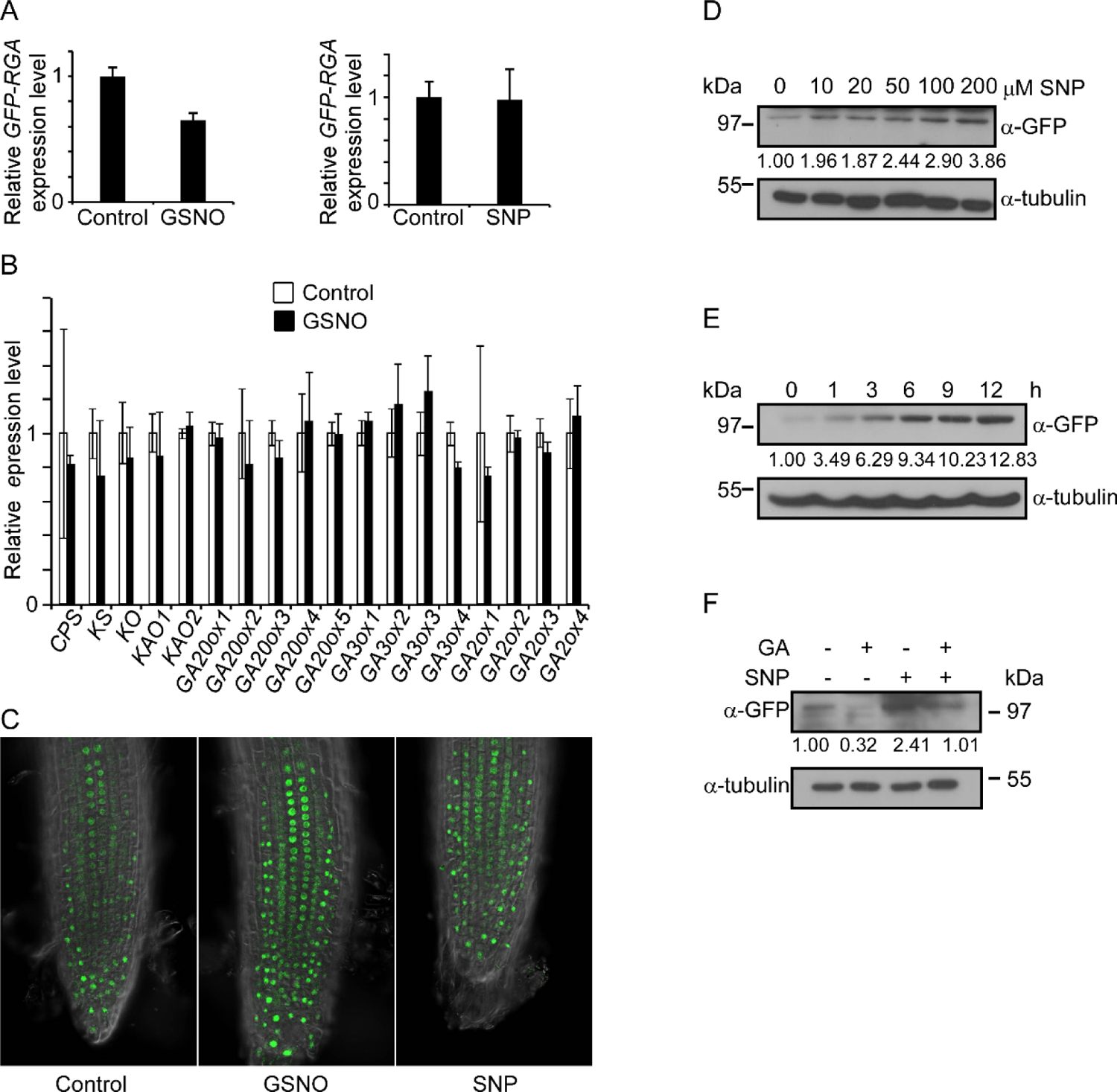
NO inhibits GA-promoted RGA degradation, related to Figure 2. (A) Analysis of the expression of *GFP-RGA* by qRT-PCR in 7-day-old *pRGA*::*GFP-RGA* transgenic plants treated with 300 μM GSNO or SNP for 6 hours. (B) Analysis of the expression of GA biosynthesis genes by qRT-PCR in 7-day-old Col-0 seedlings treated with 300 μM GSNO for 6 hours. (C) Confocal microscopic images of root tips derived from *pRGA*::*GFP-RGA* transgenic seedlings treated with 300 μM GSNO or SNP. (D and E) Immunoblotting analysis of GFP-RGA proteins in 7-day-old *pRGA*::*GFP-RGA* transgenic plants treated with indicated concentrations of SNP for 6 hours (D) and 300 μM SNP for indicated times (E) using an anti-GFP antibody, and immunoblotting with an anti-tubulin antibody is served as loading control. Quantification of GFP-RGA is shown below the blot. (F) Immunoblotting analysis of GFP-RGA proteins in 7-day-old *pRGA*::*GFP-RGA* transgenic plants treated with or without 300 μM SNP or 0.5 μM GA for 6 hours using an anti-GFP antibody. Quantification of GFP-RGA is shown below the blot.

**Figure S4.**
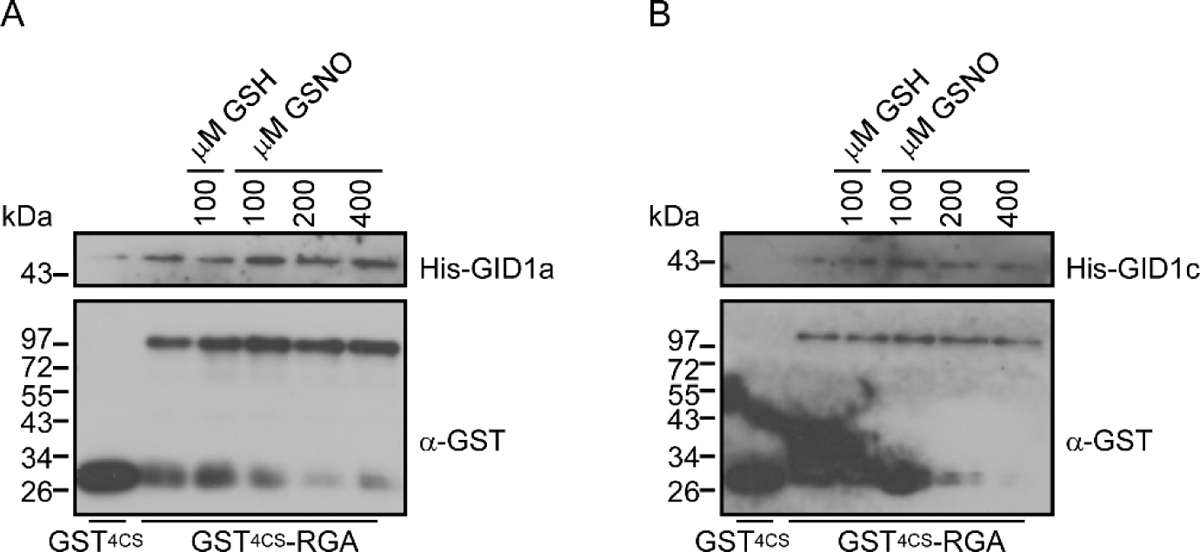
RGA-GID1 interaction is not regulated by NO, related to Figure 2. (A and B) Analysis of the interaction of His-GID1a (A), His-GID1c (B) with GST^4CS^-RGA recombinant proteins with a GST pull-down assay. GST^4CS^-RGA protein was treated with indicated concentrations of GSNO or GSH before incubated with His-GID1a or His-GID1c.

**Figure S5.**
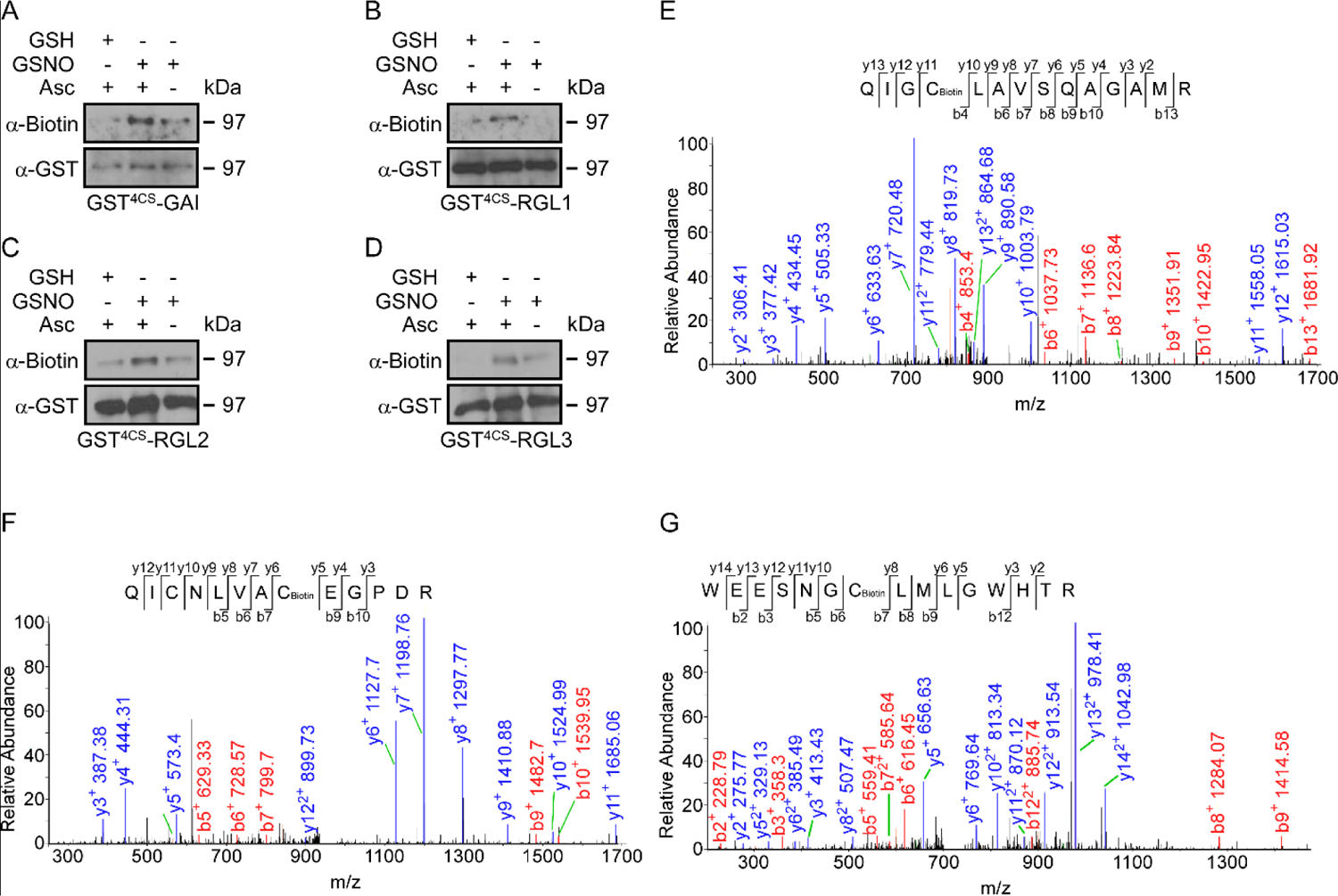
DELLA proteins are *S*-nitrosylated in vitro, related to Figure 3. (A to D) Analysis of *S*-nitrosylated GST^4CS^-GAI (A), GST^4CS^-RGL1 (B), GST^4CS^-RGL2 (C), and GST^4CS^-RGL3 (D) recombinant proteins treated with GSNO by an in vitro *S*-nitrosylation assay. Treatment with GSH and without sodium ascorbate (Asc) are served as negative controls. (E to G) Liquid chromatography tandem-mass (LC-MS/MS) spectrum of trypsin-digested and biotin-charged RGA peptides. The b- and y-type product ions are indicated, which identified Cys-249 (E), Cys-506 (F), and Cys-564 (G) as an *S*-nitrosylated residues.

**Table S1.**
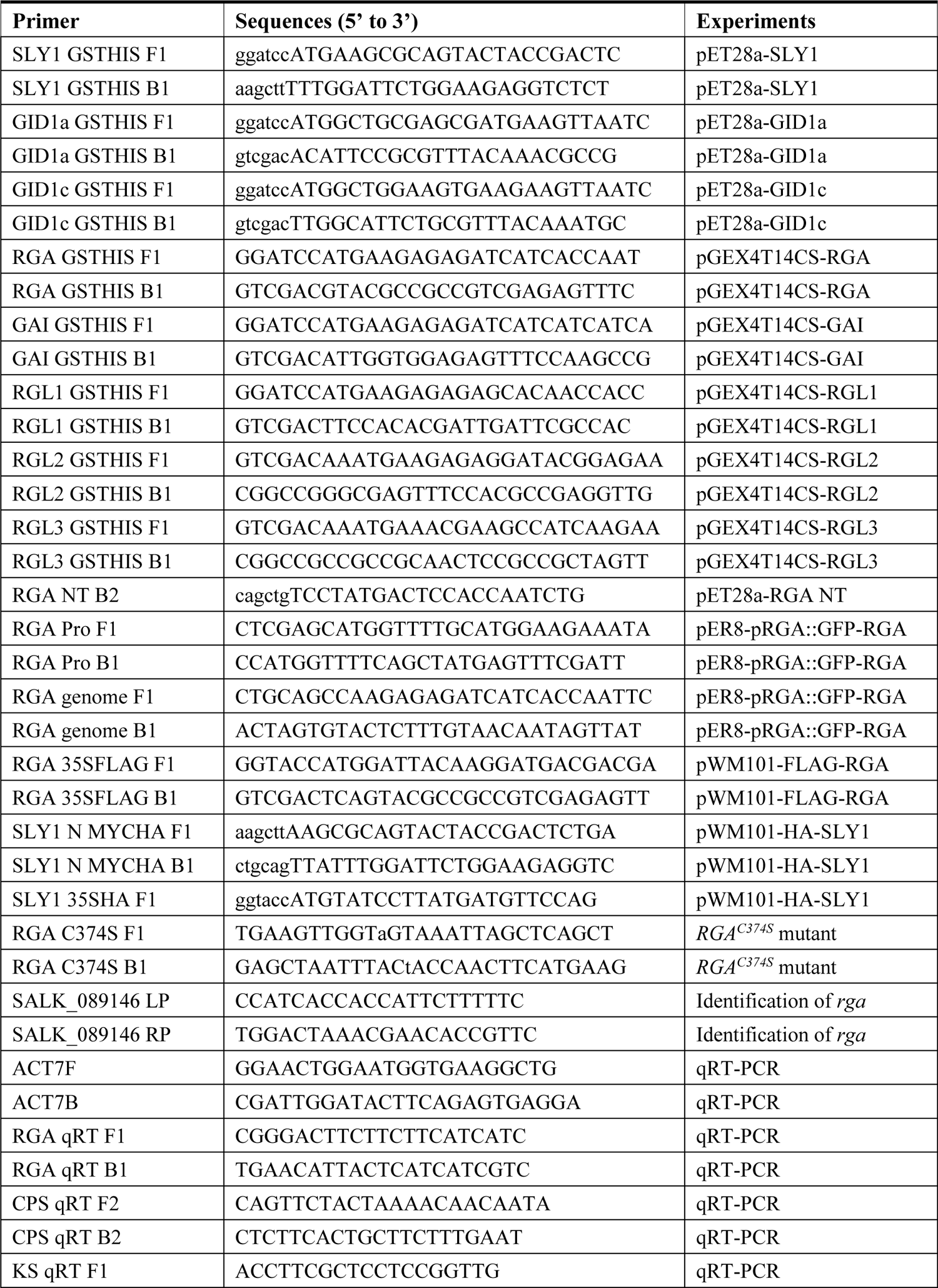
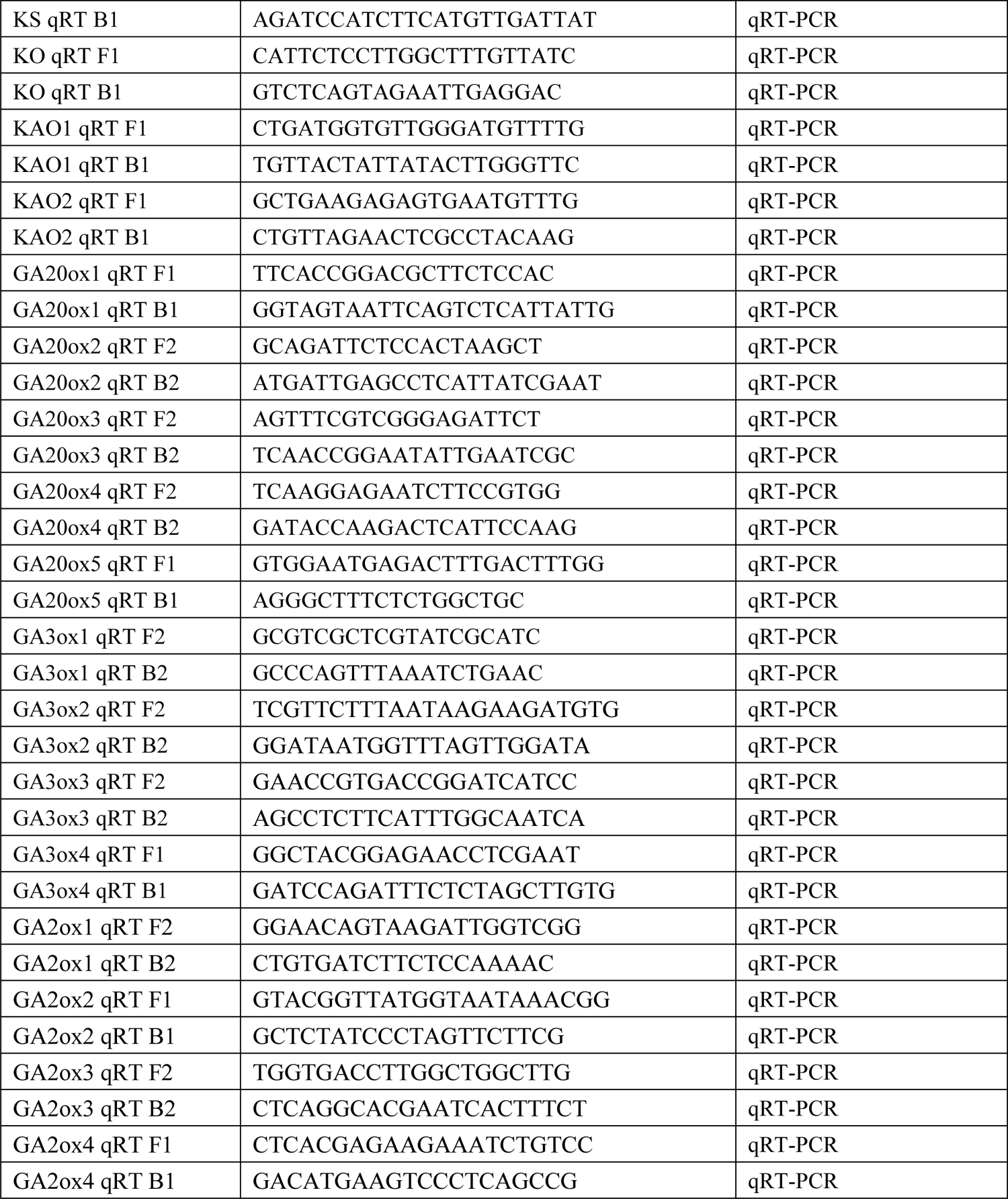
Primers used in this study, related to Figures 2, 3 and 4.

**Table S2.** *S*-nitrosylated residues of RGA, related to Figure 3

| *S*-nitrosylated residues identified in<br>mass spectrometry | Repeat 1 | Repeat 2 |
| --- | --- | --- |
| Cys-249 | √ | √ |
| Cys-374 | √ | √ |
| Cys-506 | √ |  |
| Cys-564 | √ | √ |

